# Magnetic manipulation of axonal endosome transport in live neurons

**DOI:** 10.1101/733253

**Authors:** Praveen D. Chowdary, Allister F. McGuire, Yujeong Lee, Daphne L. Che, Lindsey Hanson, Yasuko Osakada, Chinchun Ooi, Chong Xie, Shan Wang, Bianxiao Cui

**Author notes:** Corresponding Author: Bianxiao Cui, Department of Chemistry, Stanford University, 380 Roth Way, Stanford, CA 94305, USA.

## Abstract

Noninvasive control of axonal cargos in live neurons is a challenging prospect that can enable novel research on the mechanisms of axonal cargo transport, cargo-mediated signaling and axonal traffic jams in neurons. However, conventional techniques for force manipulation such as optical traps are limited to a few micron-sized cargos and are not applicable to the small axonal cargos in live neurons. Here, we present a new methodology that permits the external control of axonal endosome transport via tailored magnetic forces. By culturing neurons in a microfluidic device made up of microfabricated magnetic arrays, we can exert 3 – 48 pN forces on retrograde axonal endosomes carrying fluorescent magnetic nanoparticles, 100 – 260 nm in size. The magnetic force counters the forces exerted by molecular motors driving the endosomes and results in a wide range of perturbations on endosome transport in axons. These perturbations, captured by oblique illumination fluorescence imaging, reveal new insights on the collective function of dyneins and the nature of paused and stationary states during retrograde endosome transport in axons. Most notably, we demonstrate controllable capture and release of retrograde endosomes in axons by toggling the external magnetic field. This technical advance has great potential to elucidate the spatiotemporal origins of long-distance endosome signaling pathways as well as the ramifications of axonal traffic jams in neurons.

## Introduction

The axon acts as a conduit for long-distance cargo transport between the cell body and the axon terminals in neurons^1,2^. Axonal microtubules serve as polarized tracks for the molecular motor proteins, kinesins and dyneins, transporting in opposite directions and carrying a diverse range of cargos across the entire axon^3,4^. Numerous cargos including cytoskeletal structures, proteins, synaptic vesicles, endosomes, etc., are transported anterogradely from the cell body to the axon terminals by kinesins. The fast anterograde transport is critical for axonal growth, synaptic assembly, and plasticity in neurons. On the other hand, various cargos including vesicles, signaling endosomes, lysosomes, autophagosomes, etc., are transported retrogradely from the axon terminals to the cell body by dyneins. The fast retrograde transport mediates many signaling, clearance, and degradation mechanisms in neurons^5,6^. The long-distance axonal cargo transport is therefore fundamental to the structure, function and survival of neurons. Indeed, impaired axonal transport caused by defects in motor function or adverse conditions like traffic jams is implicated in many neurodegenerative diseases^5,7-17^.

The interplay between long-distance axonal trafficking of cargos, cargo-associated signaling mechanisms, and neuronal function, has been the subject of extensive research so far^6^. However, several questions still abound on the biophysical mechanisms of motor transport in axons, the spatiotemporal origins of signaling cargoes, and the ramifications of axonal traffic jams for neurodegeneration. A variety of such questions potentially can be addressed by the development of new technologies that enable noninvasive manipulation of axonal cargos by external forces. Firstly, the analysis of axonal cargo transport under controlled external forces can elucidate the transport machinery and cooperative mechanics of motors driving this long-distance transport. Secondly, external control of axonal cargos like signaling endosomes that mediate the retrograde signaling pathways can unravel the spatiotemporal nature of endosome signaling in different neuronal compartments. Thirdly, noninvasive force control of axonal cargos can lead to new innovative assays for engineering axonal traffic jams to systematically analyze the links between traffic jams and neurodegeneration.

However, noninvasive external force control of axonal cargos in live neurons is a technical challenge. The use of optical force in cells is limited to a few large cellular cargos such as lipid droplets and latex bead phagosomes that can be manipulated by optical tweezers^18-21^. The small axonal cargos in narrow caliber axons (0.5-1µm) are not amenable to optical trapping approaches, which are also quite invasive for live neurons. Similarly, magnetic trapping approaches^22,23^ that use noninvasive magnetic forces to control micron-sized magnetic beads have neither been used for manipulating cargo transport in cells nor can be applicable to axonal cargo manipulation. The application of microfabricated magnetic tweezers to control the diffusive accumulation of microinjected magnetic nanoparticles in cells was reported^24,25^. However, there is no reported method for targeting magnetic nanoparticles to cellular cargos. Further, none of these approaches was applied to control directional cargo trafficking driven by molecular motors in cells. Steketee and coworkers reported a novel approach using a sharp magnetic tip for endosome transport manipulation in neurons^26,27^. However, the single-tip-based approach has low-throughput (one neuron at a time) and lacks the selectivity of retrograde versus anterograde endosomes, which limit its biological applications. New innovative approaches are required for the external force manipulation of axonal cargos in live neurons.

In this work, we present a novel methodology based on microfluidic neuron culture within microfabricated magnetic arrays. Specifically, we demonstrate the use of magnetic forces to manipulate retrograde axonal endosome transport in live neurons. By selectively targeting magnetic nanoparticles (MNPs) into retrograde axonal endosomes, we could exert magnetic forces in the range of 3 – 48 pN that lead to a wide range of perturbations on MNP-endosomes in axons. Besides providing new insights on the mechanics of retrograde endosome transport in axons, our results prove that noninvasive external forces can be engineered to control axonal cargos. Our methodology, which can be optimized further by customizing the device’s design and MNPs, holds great promise to enable new research on the mechanisms of axonal cargo transport, cargo-mediated signaling and axonal traffic jams in neurons.

## Results

### Magnetic manipulation of axonal endosomes

Our approach is to induce receptor-mediated endocytosis of fluorescent magnetic nanoparticles (MNPs) at the axonal terminals, and then apply highly-localized external magnetic forces to manipulate the MNP-containing endosomes in the middle segments of axons (Fig. 1A). The basic principle is that a MNP in a magnetic field gradient is subjected to a force proportional to the field gradient and the particle’s magnetic moment^22,23^. In order to counter the cumulative force of dyneins driving the retrograde MNP-endosomes, the magnetic force must be in the range of 5 – 20 pN. The narrow caliber of axons (< 1 µm) and the small size of axonal endosomes (< 300 nm)^28,29^ limit the diameter of MNPs to be < 250 nm for effective endocytosis and retrograde axonal transport of MNPs. The magnetic field gradient required to exert 5 – 20 pN on such small MNPs is ∼10^5^ T/m based on previous measurements^30^. Such high gradients can be achieved by placing magnetic microstructures in close proximity (<10 µm away) to the axon containing the MNP-endosomes. Further, by using soft magnetic microstructures, which can be magnetized by an external magnetic field, the magnetic forces on MNP-endosomes can be toggled on/off as needed. The effect of magnetic forces on the transport of MNP-endosome can be visualized in real time by oblique illumination fluorescence imaging, which we previously demonstrated^31-33^.

**Figure 1:**
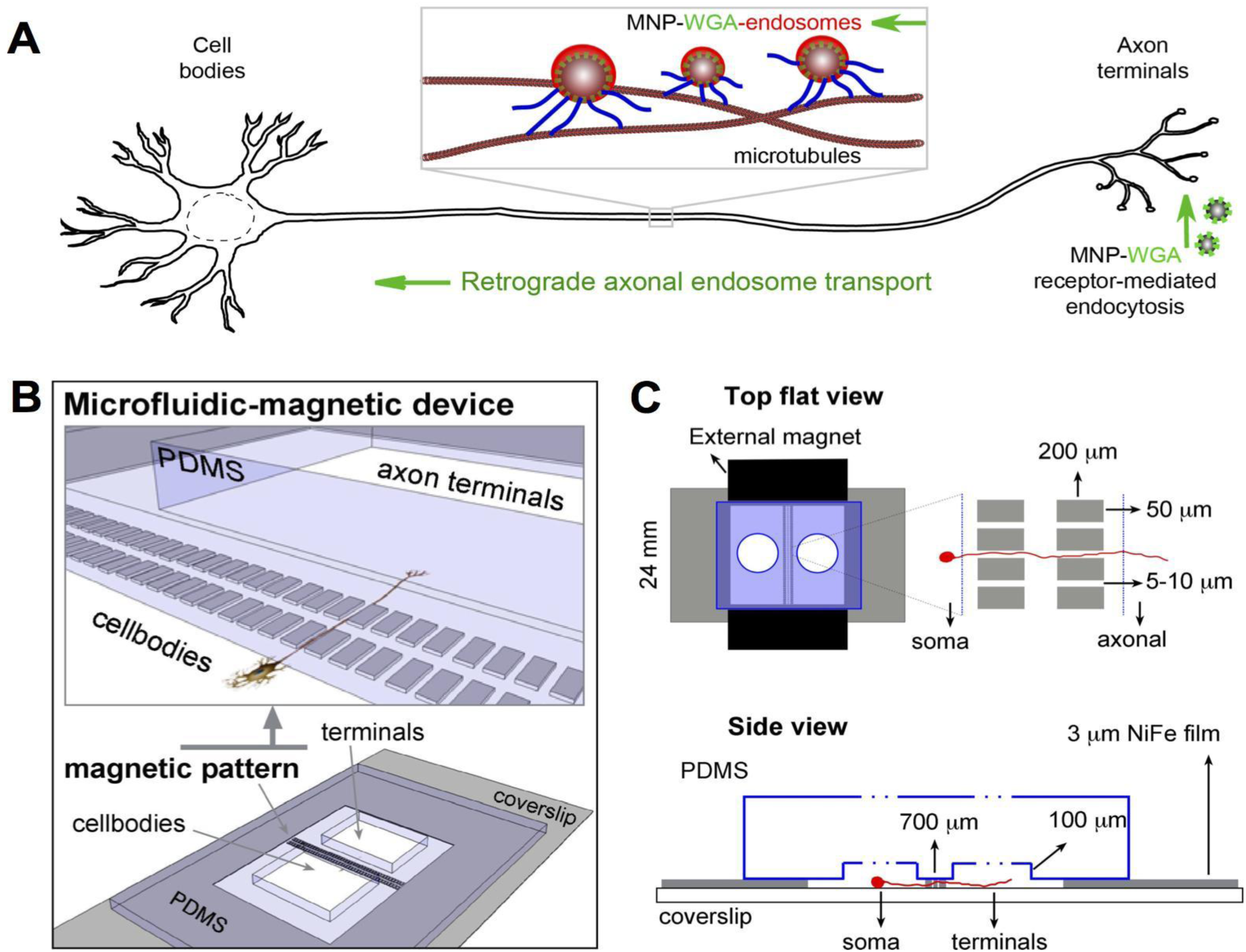
Magnetic manipulation of retrograde axonal transport A) A schematic illustration of wheat germ agglutinin (WGA)-coated MNPs internalized at axon terminals and retrogradely transported along the axon by microtubular motor transport. B) Schematic of our magnetic-microfluidic device segregating the cell bodies and axon terminals into two different chambers connected only by the channels between the rectangular micromagnets. C) Schematic of the magnetic pattern geometry and the application of external magnetic field to magnetize the micromagnets.

Therefore, the key design considerations in our methodology are A) surface functionalization of fluorescent MNPs for receptor-mediated endocytosis in neurons and imaging, B) microfluidic isolation of cell bodies and axon terminals so that MNPs can selectively be applied to the axon terminals, C) soft micromagnets that can be toggled on/off, placed in close proximity to the axons carrying retrograde MNP-endosomes, and D) compatibility with fluorescence imaging to monitor the effect of magnetic forces on axonal MNP-endosome transport in real time. We achieved all these requirements by fabricating arrays of soft micromagnets on glass coverslips and guiding axonal growth through the tight spacing (5 – 10 µm) between the micromagnets as elaborated below.

### Design and fabrication of magnetic-microfluidic device for neuron culture

Fig 1B-C shows the design of our prototype magnetic-microfluidic device that is compatible with microfluidic neuron culture, high-gradient magnetic manipulation and fluorescence imaging. The microfluidic-magnetic device features arrays of rectangular micromagnets, fabricated by patterned electrodeposition of permalloy (80:20 Ni:Fe) on glass coverslips. We chose permalloy for its excellent soft-magnetic properties. The rectangular micromagnets, passivated by a 100 nm layer of SiO_2_, are closely spaced to create 200 µm long, 5-10 µm wide, and 3 µm deep microfluidic channels on the glass surface. To achieve high throughput, hundreds of magnetic microstructures are fabricated on the same substrate. A polydimethylsiloxane (PDMS) chip with two biopsy-punched holes forms a watertight seal on the micromagnet surface, thus creating the microfluidic-magnetic device with two chambers connected by the microfluidic channels between micromagnets. Neurons plated in the cell body chamber extend their axons through the microfluidic channels into the distal axon chamber, which ensures the close proximity of axons to the micromagnets. MNPs incubated in the distal axon chamber are internalized, forming MNP-endosomes, which are retrogradely transported along the axons between the micromagnets. By applying an external magnetic field using two permanent NdFeB magnets placed on either side of the device, we can magnetize the soft micromagnets and induce the magnetic forces, which act inward into the microfluidic channels (see below). The effect of magnetic forces on MNP-endosomes within the channels is visualized in real time using oblique illumination fluorescence imaging through the transparent microfluidic channels.

The fabrication process of the microfluidic-magnetic device is illustrated step-wise in Fig. 2A. (1) The process begins with a clean standard microscope glass coverslip (24 mm x 40 mm, 120 µm thick). (2) A 10 nm/50 nm Cr/Au conducting seed layer is sputtered on the glass coverslip. (3) A 7 µm thick photoresist layer is spin-coated onto the Au layer. (4) The photoresist layer is then patterned by UV exposure in a contact mask aligner to expose the Au layer in desired areas. (5) Electroplating of permalloy (Ni:Fe 80:20, 3µm thickness) onto the exposed Au layer is carried out according to a published protocol^34^. (6) The photoresist layer is removed by washing in acetone. (7) The Cr/Au layer is removed by dry etching, which renders the coverslip transparent wherever the magnetic material is not plated. (8) The magnet surface is then passivated with a thin layer of 50nm/50nm Si_3_N_4_/SiO_2_. (9) The SMD is formed by sealing the micromagnet surface with a PDMS chip.

**Figure 2:**
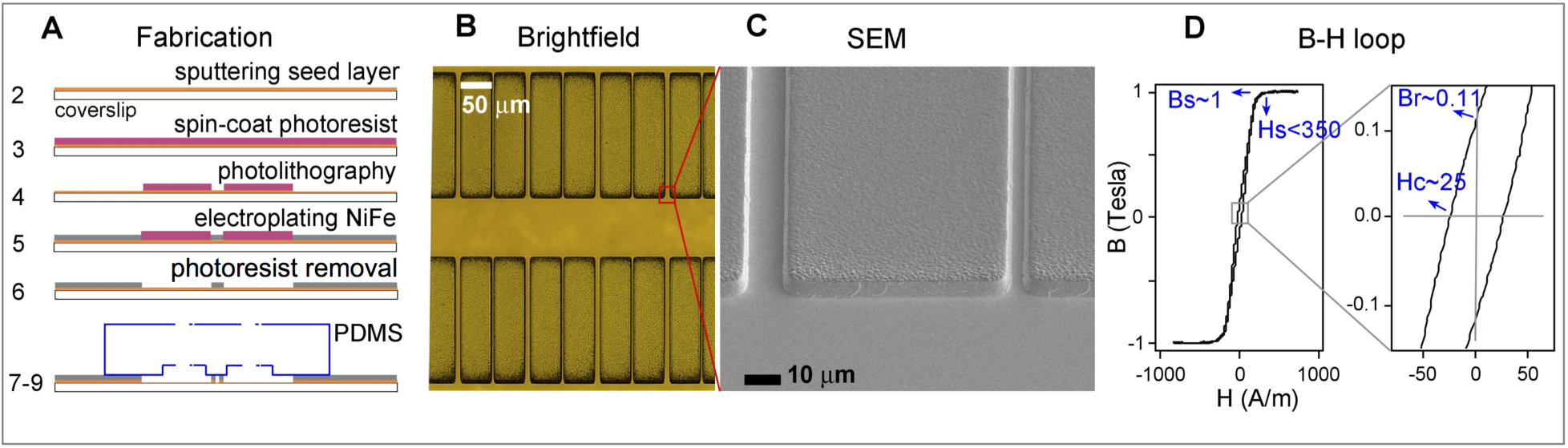
Fabrication and characterization of the magnetic-microfluidic device. A) The fabrication procedure of the micromagnets. B) Brightfield and C) SEM images of the micromagnet pattern. D) B-H hysteresis loop of the magnetic-microfluidic device shows very little remanent magnetization.

Figs. 2B,C show the brightfield and scanning electron microscopy (SEM) images, respectively, of the micromagnetic pattern. The rectangular micromagnets are alternately spaced by 5 µm and 10 µm distances. We note that the external NdFeB magnetic field is applied along the width of the rectangular micromagnets to magnetize them and induce the poles along their width. The shape anisotropy of the rectangular micromagnets ensures that the micromagnets demagnetize almost completely when the NdFeB field is removed. Fig. 2D shows the hysteresis loop of the microfluidic-magnetic device measured using a B-H loop tracer (Shb Instruments, Inc.). This confirms the soft magnetic behavior of the permalloy film with a Bs value of 1 T at an applied field of 350 A/m, low remanance of 0.11 T and a coercivity of 25 A/m. The low remanence shows that we can turn these micromagnets on/off by using an external NdFeB magnetic field.

### Magnetic force directionality and force estimates within SMDs

In order to analyze the magnetic forces in our SMD (softmagnetic-microfluidic device), we first simulated the magnetic field gradients in the channels between micromagnets using a 3D analytical model^35,36^. Magnetic field gradients as high as 10^5^ T/m are calculated within 1 µm from the micromagnet edges (Fig. S1, Supporting Material). Using a linear saturation magnetization model^35^, we then estimated the magnetic forces to be in the range 5 – 50 pN on a 100 nm iron oxide MNP within our microfluidic-magnetic device (Fig. S2). A notable feature of the magnetic pattern is the directionality of magnetic force within and outside the channels, as shown by the results of simulations in Fig. 3A. The magnetic force within the channel primarily acts perpendicularly to the channel (i.e., to the axonal transport direction). At the channel exits, however, the force is directed inward along the channel (i.e., parallel to the transport direction). MNP-endosomes transporting out of the channels would therefore face an opposing magnetic force acting as a roadblock. This directionality of magnetic forces is also confirmed by experimentally tracking the MNPs under force in 10:1 glycerol:water mixtures on our SMDs (Movie S1). Fig. 3B shows the time-projection of a movie depicting the tracks of MNPs pulled towards the micromagnets when the forces are turned on.

**Figure 3:**
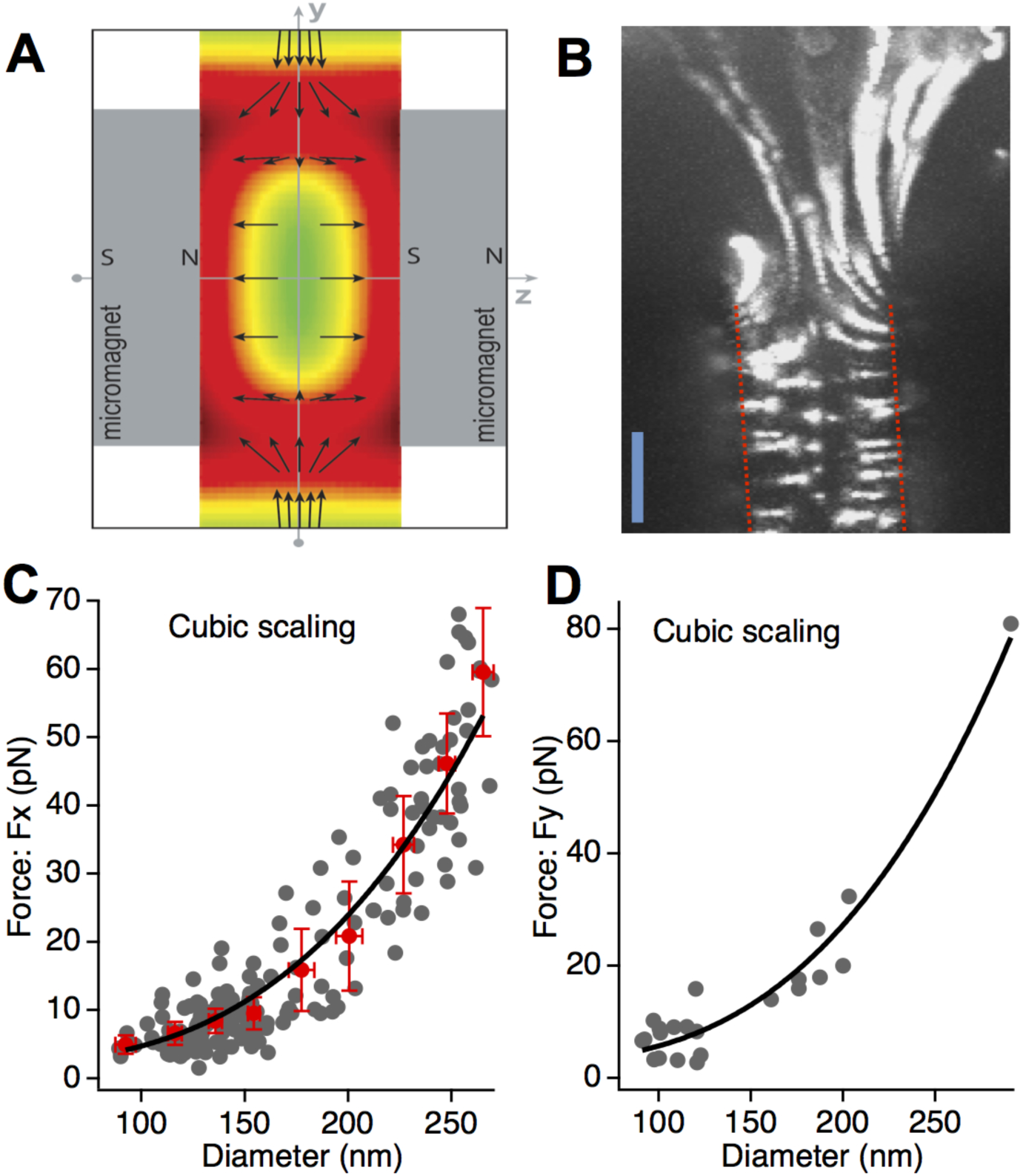
Magnetic force estimation in the magnetic-microfluidic device A) Directionality of the magnetic force structure within the channel between two micromagnets. The color profile highlights regions of high force (red) and low force (green). B) Trajectories of fluorescent MNPs being pulled towards the micromagnetic edges (red dotted lines) under force. Scale bar = 5 μm. C) *F*_*x*_ at a distance of 2.5 μm from magnet edge within the channel as a function of particle diameter. Individual data points from each MNP trajectory (gray markers) and binned statistics (Mean±3*SEM in red). D) *F*_*Y*_ at a distance of ∼3 μm from the channel opening as a function of particle diameter. Binned statistics were not computed due to limited data above 200nm.

We also experimentally estimated the magnetic forces on MNPs within and outside the channels from the MNP trajectories under force in 10:1 glycerol:water mixtures (Supporting Material). First and foremost, we observed a clear mapping between the MNP size (SEM diameter, *D*) and MNP-fluorescence intensity (*I α D*^*2*^) as shown by the overlapping distributions of *D* and ***√I*** in Fig. S3. This enabled us to estimate a MNP’s size from its fluorescence intensity (***D = A*√I+B***) by matching the cumulative distribution functions (Fig. S3). Using the MNP size *D* and the MNP velocity *v*, extracted from individual MNP trajectories under force, viscosity *η*, we estimated the force (*F=3πηvD*) on MNPs within and outside the channels at different distances from the micromagnets (Fig. S4). Fig. 3C shows the magnetic force on MNPs of different sizes at a distance of 2.5µm from the micromagnet edge within the channel. Fig. 3D shows the magnetic force acting inward to the channel at ∼ 3 µm distance from the channel exits. The near-cubic scaling of magnetic force with MNP size further validates the mapping between the MNP fluorescence intensity and MNP size (Figs. 3C,D). In what follows we used these cubic scaling curves to estimate the force on an MNP in our microfluidic-magnetic device, given the measured fluorescence intensity of the MNP.

### *Axonal transport of MNPs in* microfluidic-magnetic device

We tested the compatibility of microfluidic-magnetic devices for long-term neuronal culture using a protocol of culturing DRG neurons in microfluidic devices from our earlier studies^31,32^. Briefly, the SMD is coated with poly-L-lysine to facilitate cell attachment and survival. DRG neurons from embryonic E18 rats are dissociated and plated into the cell body chamber. Within 3 – 5 days, the neurons readily extended the axons across the two arrays of micromagnets into the distal axon chamber. DRG neurons cultured in the microfluidic-magnetic devices survived > 3 weeks, which is comparable to neuronal survival on poly-L-lysine coated coverslips in our earlier studies. Fig. 4A shows the brightfield image of a 12^th^ day DRG culture in a microfluidic-magnetic device. The shade in the cell body chamber (left side of the image) is due to the uneven edge of the manually-cut PDMS chip. The tiny black dots in the distal axon chamber are the lectin-coated MNPs (see below) applied for the imaging of retrograde MNP-endosome transport. The micromagnet pattern is quite durable and the microfluidic-magnetic device can be reused for multiple culturing, imaging, and cleaning cycles.

**Figure 4:**
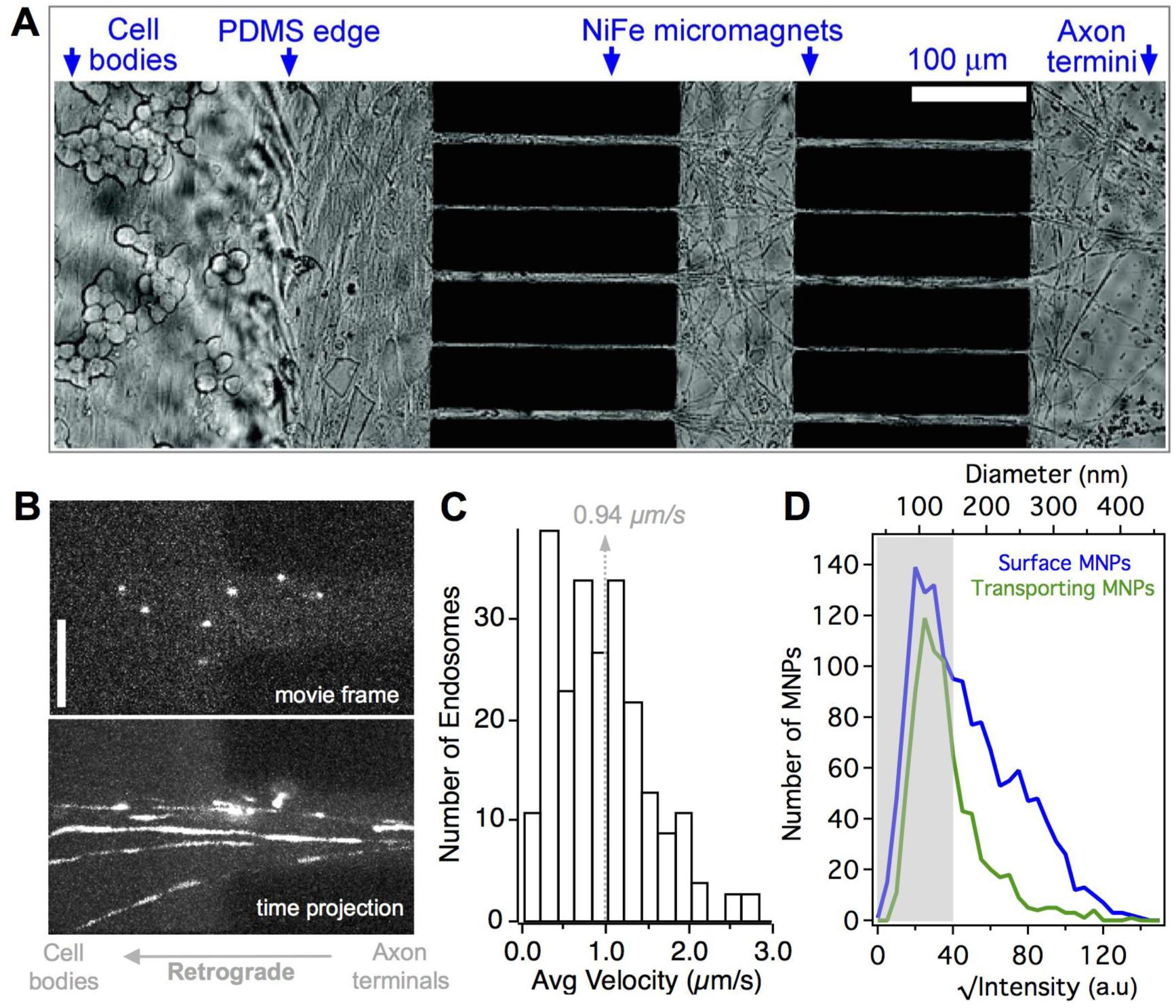
Retrograde axonal transport of MNP-endosomes in magnetic-microfluidic device A) DRG neuron culture in a magnetic-microfluidic device at 12 days in vitro shows many axons crossing the magnetic channels into the distal axon compartment. B) Snapshot of a movie showing MNP-endosomes transporting in axons (top) and the time projection of the movie showing the MNP-endosome tracks along the axons (bottom). Scalebar = 10 μm. C) Average speed of MNP-endosomes retrogradely transporting in axons. D) Fluorescence intensity distributions of surface bound MNPs (blue) and MNP-endosomes transporting in axons (green). The corresponding MNP diameter shown on top axis is obtained from the mapping ***D = A*√I+B***.

We then studied the retrograde transport of wheat germ agglutinin (WGA)-coated fluorescent MNPs (100 – 350 nm, SEM size distribution in Fig. S3) in axons using oblique illumination imaging with minimal laser power (∼0.14 W/cm^2^). WGA is an established retrograde tracer^37,38^ that is shown to promote receptor-mediated endocytosis and transport of nanoparticles in cells^39^ including neurons^31^. We observed robust retrograde transport of WGA-MNPs in the axons within microchannels after 2.5 h of incubation in the distal axon chamber of the SMD (Movie S2). Fig. 4B shows a snapshot of MNP-endosomes in axons and the projection of a time-lapse movie of MNP-endosomes moving out of the micromagnetic channel into the cell body chamber (magnetic force off). Fig. 4C shows the speed distribution of the retrograde MNP-endosomes in axons with an average moving speed at 0.94 ± 0.55 µm/s, which agrees with previously published work for axonal transport studies^31,32^. Fig. 4D shows the fluorescence intensity distribution of the retrograde MNP-endosomes transporting in axons superimposed on that of the incubated MNPs. Based on the mapping between MNP’s fluorescence intensity and size, there is robust internalization and retrograde transport of MNPs <150 nm in size (shaded region in Fig 4D). We also see the retrograde transport of MNPs up to a size range of 260 nm, although with a decreased propensity than that of smaller sized particles.

### Effect of magnetic force on MNP-endosome transport in axons

When the external NdFeB magnets are introduced to magnetize the microfluidic-magnetic device, the resulting forces lead to a wide range of perturbations on the MNP-endosomes transporting in the axons. The 100 – 260 nm size range of MNPs being transported by the endosomes approximately corresponds to magnetic forces in the range of 3 – 48 pN for axons within 2.5 µm of the edge of micromagnets (Figs. 3C,D). While the transport of small MNP-endosomes (100 nm, ∼3 ± 1.7pN) force is minimally perturbed by the magnetic forces, the large MNP-endosomes (260 nm ∼ 48 ± 7.5pN) are stalled and trapped on the edges of micromagnets. The trapped endosomes are released and resume normal retrograde transport when the magnetic force is turned off by removing the external NdFeB magnets. In what follows, we present these results and the insights gleaned on the mechanics of retrograde endosome transport in axons.

### Catch and release of retrograde MNP-endosomes within microchannels

Fig. 5A is the time projection of a time-lapse movie showing a retrograde MNP-endosome transporting along the axon with the magnetic forces turned on. We estimate a magnetic force of ∼46 ± 7.5pN acting on the MNP-endosome (from the MNP size ∼255 nm, based on intensity) and the distance from the micromagnet (<2 µm). Interestingly, the stably moving MNP-endosome exhibited some back-and-forth oscillations before being pulled and trapped at the edge of the magnet (Fig. 5B, Movie S3).

**Figure 5:**
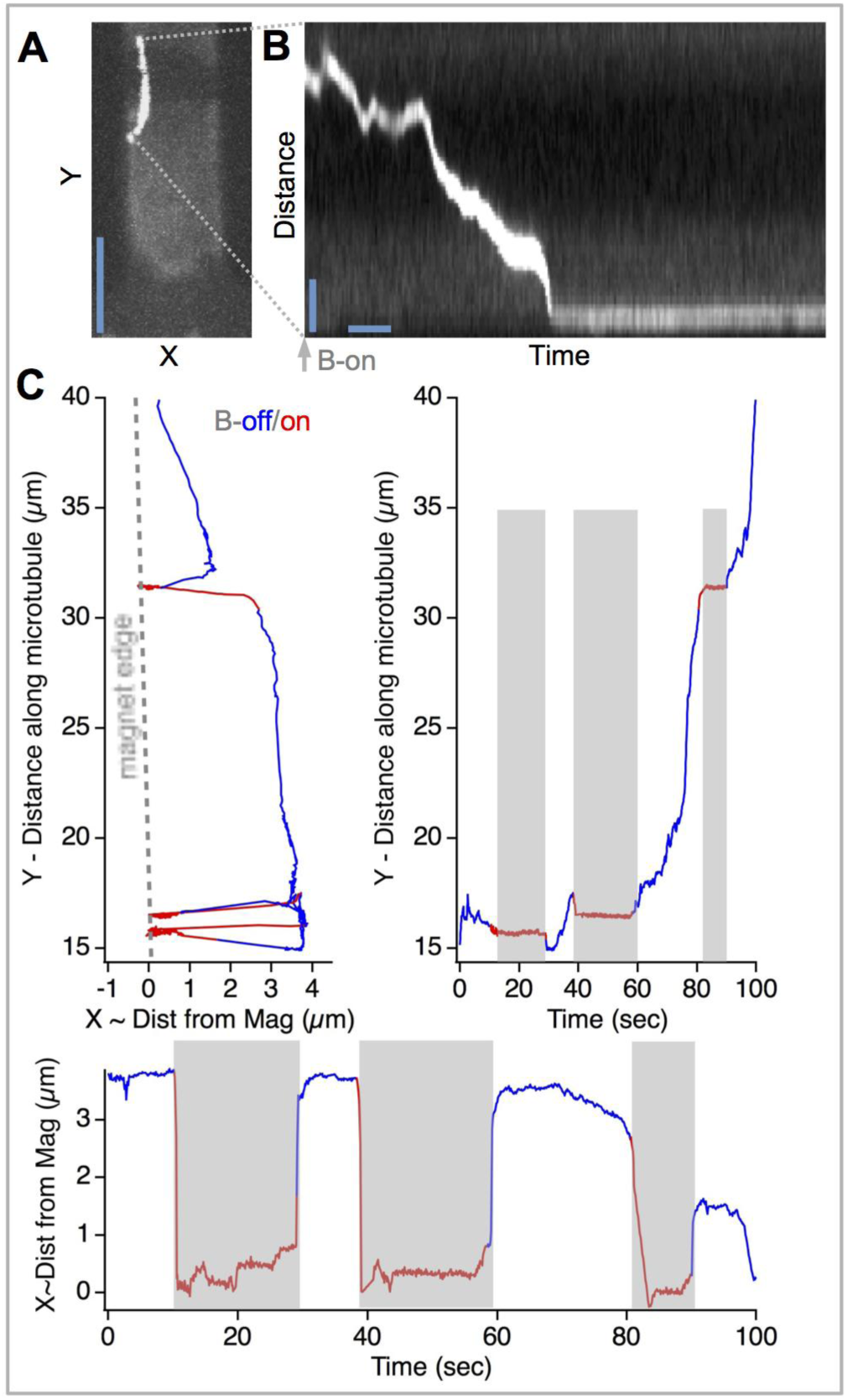
Magnetic force perpendicular to the direction of MNP-endosome transport is sufficient to stall the transport of large MNPs. A) Time projection of a movie showing the track of an MNP-endosome pulled towards the magnet edge under force. Scalebar =10 μm. B) Kymograph depicting the trajectory of the MNP-endosome in A. Scalebars = 2 μm, 3 s C) Trajectory of a retrograde MNP-endosome repeatedly captured and released by toggling the magnetic field on and off. Y versus X (top left, the dashed line represents the edge of micromagnet), Y versus time (top right) and X versus time (bottom) trajectories are colored in red when the magnetic field is on. The X coordinate is referenced with respect to the magnet edge. The shaded regions correspond to when the external magnetic field B (and thereby the magnetic force) is on.

In many cases with such large MNP size, the magnetic perturbation is quite severe. Fig. 5C, shows the time projection of a movie with a retrograde MNP-endosome (MNP size ∼259 nm) transporting along the axon when the magnetic force is toggled on and off for three cycles (Movie S4). When the magnetic forces are suddenly turned on (trajectory segments colored in red in Fig. 5C), the MNP-endosome is pulled close to the magnet edge, possibly deforming the local axonal shape within the channel or displacing the axon towards the magnet. When the magnetic force is turned off, the MNP-endosome recoiled back to its original position and resumed robust unidirectional retrograde transport indicating that the magnetic force does not affect the endosome membrane or the local microtubule structure. In as much as the entire axon might have moved or deformed towards the magnet edge we cannot corroborate if the MNP-endosome detached from the microtubule in this case. This capture and release of the MNP-endosome between processive retrograde runs could be repeated multiple times (Movie S4). These results indicate that by using MNPs (∼250 nm) in our prototype SMDs, we could effectively arrest and release retrograde MNP-endosome trafficking in the axonal segments within the microchannels.

Similar capture and release events were seen even for a few smaller MNP-endosomes (∼150 nm MNP size) when the axon is within 1.5 µm of the edge of micromagnet (Movie S5). However, in general the MNP-endosome transport with MNP size < 150nm (∼9 ± 2.4pN) is minimally perturbed within the microchannels. Surprisingly, we also observe many endosomes with MNP size ∼200 nm transporting without being stalled within the microchannels despite the high estimated force of ∼22 ± 7.5pN. In order to rationalize this, it is important to note that the force on MNP-endosomes within microchannels (especially in axons >2µm from the edge of magnets) is orthogonal to the endosome transport direction. It is plausible that this orthogonal force is buffered by the endosome pressing against the axonal cytoskeleton pressing against the endosome depending on the proximity between the axonal membrane and the MNP-endosome. Therefore, the actual projected load on the MNP-endosome could be smaller and countered by the cooperative activity of multiple dyneins. When the orthogonal force is strong enough to pull the microtubule-bound MNP-endosome to the edge of the magnet (possibly displacing the axon), a significant force component results which opposes transport thereby trapping the endosome. Our data suggests that this is the case for MNP-size ∼250nm with the estimated force of ∼43 ± 7.5 pN.

### Opposing magnetic force at channel exits

We then studied the effect of opposing magnetic force on MNP-endosomes by monitoring the retrograde MNP-endosomes transporting out of the microchannels in the SMDs (Movie S6). We observed a wide range of force perturbations on the MNP-endosomes at the channel exits. Endosomes with small MNPs (<150 nm) transported out of the channels with minimal perturbation (Fig. 6, in green). Many MNP-endosomes gradually moved out of the channels but generally showed a propensity to pause and undergo brief reversals at channel exits. Notably, endosomes with larger MNPs exhibited repeated stalls, detachments and long recoils under the influence of the opposing magnetic force (Movie S6, Fig. 6 in red). Some of these perturbed MNP-endosomes were gradually pulled back and remained stationary within the axons at channel exits (Fig. 6). The varying effect of magnetic forces seen on different MNP-endosomes is primarily due to the polydisperse size distribution of MNPs in this study as well as the alignment of different axons and their proximity to magnets at the channel exits. By using a narrow distribution of LARGE MNPs (∼250nm), we can potentially achieve a uniform force regime that effectively blocks the exit of MNP-endosomes from the channels in the SMDs.

**Figure 6:**
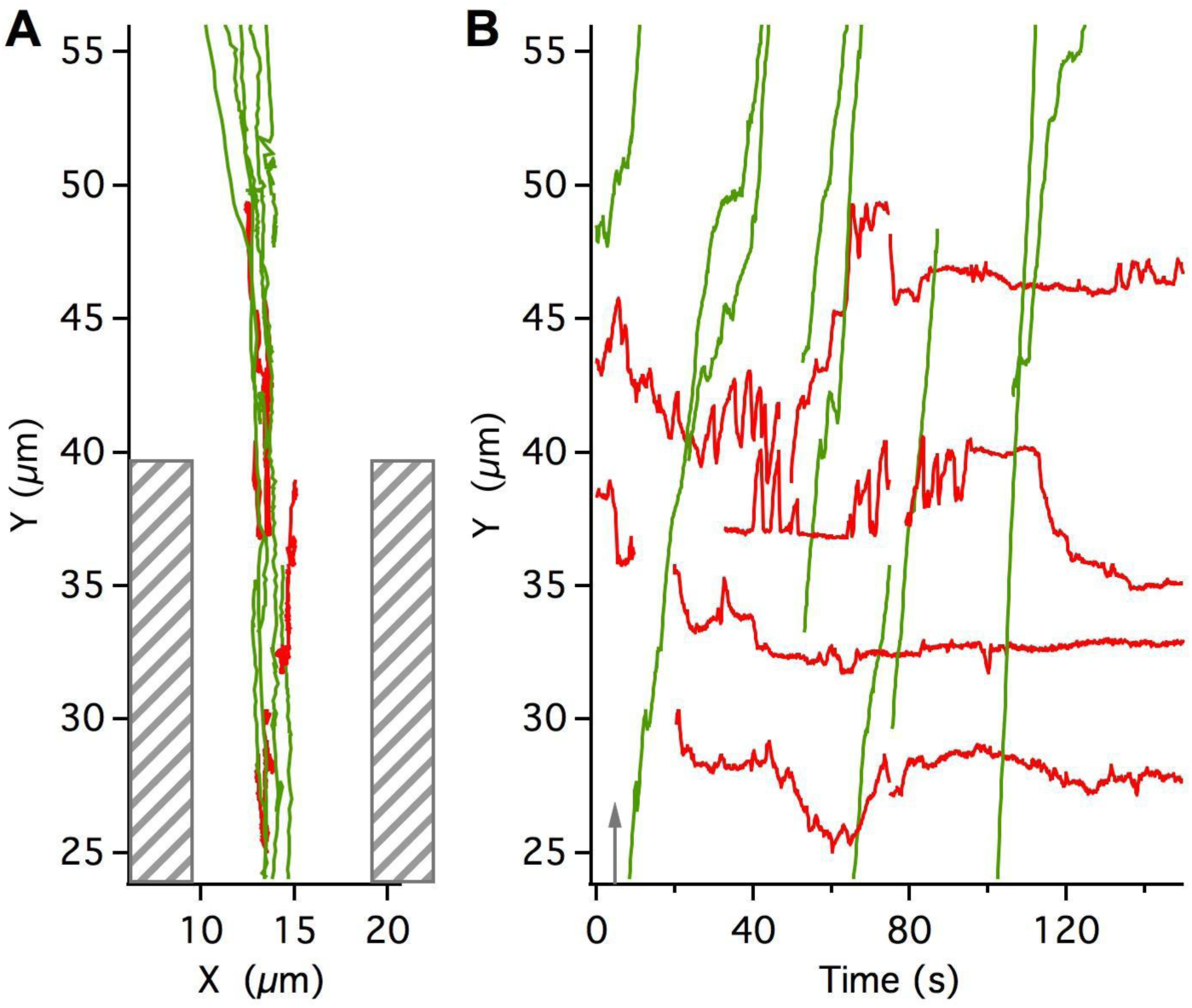
Magnetic force opposing the MNP-endosome transport at the channel exit. A) The Y-X axonal tracks of a few MNP-endosomes with respect to the micromagnet positions. B) The MNP-endosome trajectories (Y versus time), which approximately correspond to the distance traversed along the axon with time. While the endosomes with smaller MNPs are minimally perturbed (green) and are transported out of the channels, the endosomes with larger MNPs exhibit repeated detachments and are stalled around the channel exit (red). Also see Movie S4. The magnetic force is turned ON at the start (within 1-3 seconds) of the movie.

### Perpendicular force on paused retrograde endosomes - pauses are microtubule-bound states

One of the notable features of axonal endosome transport is the frequent pausing exhibited between processive runs along the microtubules. Earlier reports have shown that retrograde axonal endosomes pause for about 30% of the time during retrograde transport ^6,32^. Here, we analyzed the effect of small (2-6 pN) magnetic forces, perpendicular to the microtubule, on the paused states of retrograde MNP-endosomes in the microchannels. Fig. 7A shows an MNP-endosome trajectory transporting within a distance of 2 µm from the magnet edge. The MNP-endosome transport is minimally perturbed, as the estimated perpendicular force of 5.7 ± 1.8 pN on this MNP (129 nm) is not high enough. Interestingly, we observed no displacement of the MNP-endosome towards the magnets during pauses. The MNP-endosome position during pauses is symmetric to the direction of transport (i.e., the microtubule) as shown in Fig. 7B. If the paused endosomes were unbound from the microtubule or were loosely anchored, we would expect the endosomes to drift towards the edge of the axons closer to the magnets. Our result indicates that the paused endosomes are tightly anchored to the microtubule and are not displaced by 2-6 pN of perpendicular force. This is consistent with our earlier motion analysis, which gave an effective diffusion constant of ∼0.002 µm^2^/s for the paused retrograde endosomes in axons^32^. Another common feature of axonal vesicular transport, in general, is that a subpopulation of vesicles is stationary within axons. While the exact mechanism and the relevance of these stationary states is unknown, we see that a minor fraction of retrograde MNP-endosomes exhibits long stationary states within axons. Based on a similar analysis on the effect of perpendicular magnetic forces, we found that these stationary states of retrograde MNP-endosomes are tightly anchored within axons.

**Figure 7:**
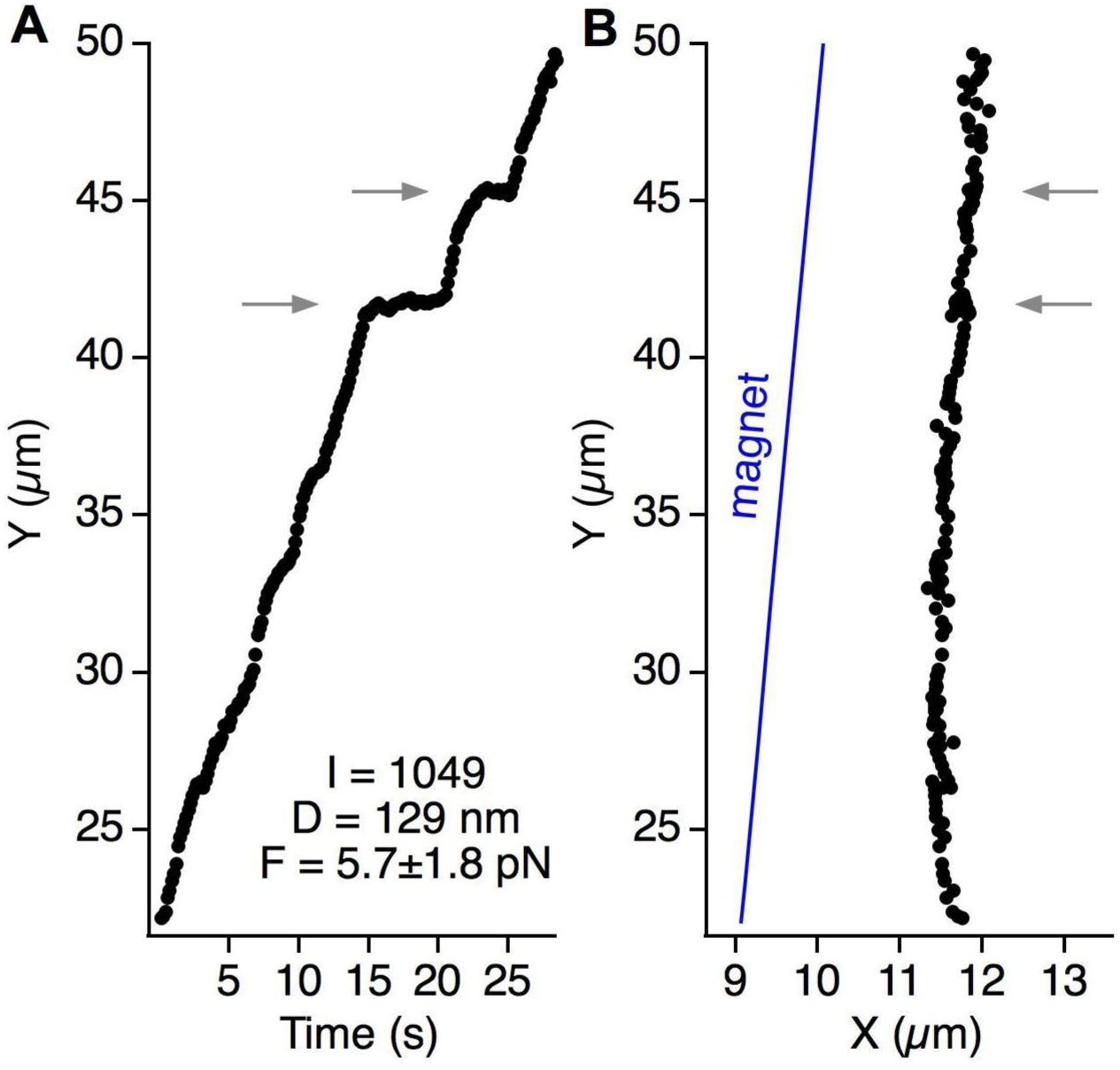
Most paused endosomes are minimally affected by the magnetic forces, indicating bound states during pauses. A) Trajectory of a retrograde MNP-endosome (Y versus time) in the presence of a perpendicular magnetic force ∼5.7 pN. B) The Y-X track of the MNP endosome is shown with respect to the edge of the micromagnet (right panel). The gray arrows point to the paused states of the MNP-endosome.

## Discussion

In this work, we report a major technical advance that permits us to manipulate axonal endosome trafficking in live neurons using noninvasive magnetic forces. We demonstrate proof of the concept that the axonal endosomes can controllably be captured and released in our prototype SMDs. Our methodology can be optimized further by more sophisticated designs of sharp-tip based micromagnet geometries and customizing the MNPs. For instance, replacing the rectangular magnets with diamond shapes with sharp tip-curvature can enhance the magnetic gradients and thereby the forces on MNP-endosomes. The choice of soft magnetic alloy material and the material for MNP are also viable improvements. Further, by using a narrow size range of MNPs as reported in literature^40^, we expect a uniform force range for more effective control on all MNP loaded endosomes in axons.

Though our focus in this work was on lectin-coated retrograde endosomes, our approach can readily be applied for the manipulation of retrograde and anterograde endosomes carrying growth factors like NGF and BDNF which mediate key signaling pathways in neurons^6,41,42^. The spatial localization of signaling endosomes within axons using noninvasive external forces is a great prospect to probe A) the functional interplay between long-distance cargo trafficking and signaling, and B) the local versus long-distance cargo-mediated signaling mechanisms in neurons. The use of sharp tip-based micromagnet geometries could also permit the controlled induction of axonal traffic jams with precise spatial control along the axons. This could open new avenues to probe the neuronal response to traffic jams and provide new insights on the causal links between impaired transport and neurodegeneration mechanisms.

## Materials and Methods

A detailed description of data processing and the experimental methods (including nanoparticle conjugation with fluorophores and WGA, oblique illumination imaging of axonal transport in soft magnetic-microfluidic devices) is provided in the Supplementary Information.

## Acknowledgements

This work is supported by the US National Institute of Health (DP2-NS082125) and the National Science Foundation (award no. 1055112 and 1344302). A.F.M. acknowledges support from the Stanford Bio-X Bowes Graduate Fellowship.

## Author Contributions

P.D.C., B.C. designed the research and P.D.C., A.F.M., Y. L. carried out the research and data analysis. A.F.M., C.X. contributed to the fabrication of magnetic devices and Y.O., D.L.C., contributed to the dissection and culturing of neurons. C.X., L.H., C.O., S.W. contributed to the device design and nanoparticle characterization. P.D.C., B.C. wrote the manuscript.

