## Supplementary material for "Magnetic manipulation of axonal endosome transport in live neurons": SupplFiguresMaterials

### Supplementary Information: Table of Contents


#### 1. Experimental methods

1.1 Materials

1.2 Conjugation of magnetic nanoparticles (MNPs) with fluorophores and WGA

1.3 Oblique illumination imaging of retrograde MNP-endosome transport in axons

#### 2. Analytical model simulations

2.1 3D analytical model for the magnetic field and gradient calculation

2.2 Magnetic forces on MNPs - Linear magnetization model:

#### 3. Supplementary data and processing methods

3.1 Mapping between MNP Size (SEM diameter) and MNP Fluorescence

3.2 Magnetic force estimation in glycerol:water mixtures

3.3 Time-lapse movie processing to generate MNP-endosome trajectories


### 1. Experimental methods

##### *1.1 Materials:*

The important chemicals used in this work and their sources are listed here. Biotin-WGA (Vector Labs). Biotin-Atto565 (Sigma Aldrich). Negative Resist NR7-6000PY (Futurrex).

Nickel chloride hexahydrate, nickel sulfate hexahydrate, iron sulfate heptahydrate, boric acid, saccharin (Electrodeposition materials, from Sigma Aldrich).

##### *1.2 Conjugation of magnetic nanoparticles (MNPs) with fluorophores and WGA:*

Superparamagnetic iron oxide magnetic nanoparticles (MNPs, 100 – 300 nm) with a high density of streptavidin surface functionalization (binding ~7000 pmol biotins per mg) were purchased from Ocean Nanotech. These MNPs were coated sequentially with fluorescent biotin-ATTO and biotin-WGA to obtain fluorescent MNP-WGA as follows. First, we incubated the MNPs and biotin-Atto565 (1:15000 MNP:dye ratio) at 4 $^{o}$C overnight in PBS with 0.05% Tween20. We then incubated the dye-coated MNPs with biotin-WGA (1:8000 MNP:WGA ratio) for 30 min before adding an excess of biotin to the occupy the residual streptavidin on MNPs. We then performed three sequential magnetic separations in a high-gradient magnetic separator (Supermag, Ocean Nanotech) to discard unconjugated biotin-dye/biotin-WGA in the reaction mixture and made up the fluorescent MNP-WGA conjugate in PBS at a final particle concentration of 0.5 – 1 nM.

##### *1.3 Oblique illumination imaging of retrograde MNP-endosome transport in axons:*

The use of microfluidic devices for primary neuron culture and imaging axonal transport had been documented by us elsewhere[1-4]. We used a similar approach for magnetic-microfluidic devices (MMDs) in this work. Briefly, an inverted microscope (Nikon Eclipse Ti-U), customized with 561nm laser excitation was set for oblique illumination. The laser beam was expanded to 3 cm diameter and focused at the back focal plane of the microscope objective (CFI Plan Apo TIRF, 100X/1.49NA) for a collimated illumination with beam waist ~120μm. Starting from total internal reflection geometry, the incident angle was gradually lowered below the critical angle till the MNP-endosomes within the axons are made visible by the oblique angle illumination. Since the microfluidic channels in our device are 3 μm high, a penetration depth of 1-3 μm is sufficient to illuminate axonal fluorophores in multiple focal planes. The fluorescence collected by the objective was relayed and focused onto an EMCCD sensor (Andor Ixon DU-897).

We used 7-12 days old DRG neuron cultures for the MNP-WGA transport and manipulation studies. The axon terminals in the MMD culture were incubated with freshly prepared MNP-WGA (0.5-1nM) for 45 min, which was then washed off by the culture medium. Shortly before imaging, the culture medium was replaced by $CO_2$-independent medium and the culture was imaged on a water-heated custom microscope stage set to maintain the culture at 37 ºC. Imaging was started typically 2.5 h after the incubation start and restricted to < 45 min session. Time-lapse movies of endosome transport were acquired at 6.7 frames per second (150ms exposure per frame) and the imaging laser power was 0.013 mW (i.e. 0.14 W/cm$^2$, for a beam waist ~120 μm).

### 2. Analytical model simulations

##### *2.1 3D Analytical model for the magnetic field and gradient calculation:*

A detailed analytical model for predicting the 2D magnetic field and the magnetic forces on nanoparticles in a magnetophoretic microsystem had been discussed in literature[5,6]. We adapted this model to our 3D microsystem in order to compute the magnetic fields and gradients within the microchannels of our magnetic-microfluidic devices (MMDs). Briefly, our device is comprised of two linear arrays of rectangular micromagnets that are magnetized by the external bias field of two NdFeB permanent magnets in the geometry shown in Fig S1A. The magnetic field at the surface of the coverslip within the microchannels between the micromagnets (shaded blue in Fig. S1A) is therefore the result of the bias magnetic field $H_b$ from the NdFeB magnets and the micromagnetic field $H_p$ from the array of permalloy micromagnets. The analytical formulae for the magnetic field B at any point (x,y,z) outside a rectangular bar magnet have been reported[6] and are given below (reference to Fig. S1B, where the polarization is along the Z-axis).

$$B_x(x,y,z) = \frac{\mu_0 M_s}{4\pi} \sum_{k=1}^{2} \sum_{m=1}^{2} (-1)^{k+m} \ln\left[ \frac{(y-y_1) + \left[(x-x_m)^2 + (y-y_1)^2 + (z-z_k)^2\right]^{1/2}}{(y-y_2) + \left[(x-x_m)^2 + (y-y_2)^2 + (z-z_k)^2\right]^{1/2}} \right]$$

$$B_y(x,y,z) = \frac{\mu_0 M_s}{4\pi} \sum_{k=1}^{2} \sum_{m=1}^{2} (-1)^{k+m} \ln\left[ \frac{(x-x_1) + \left[(x-x_1)^2 + (y-y_m)^2 + (z-z_k)^2\right]^{1/2}}{(x-x_2) + \left[(x-x_2)^2 + (y-y_m)^2 + (z-z_k)^2\right]^{1/2}} \right]$$

$$B_z(x,y,z) = \frac{\mu_0 M_s}{4\pi} \sum_{k=1}^{2} \sum_{n=1}^{2} \sum_{m=1}^{2} (-1)^{k+n+m} \tan^{-1}\left[ \frac{(x-x_n)(y-y_m)}{(z-z_k)\left[(x-x_n)^2 + (y-y_m)^2 + (z-z_k)^2\right]^{1/2}} \right]$$

Eq. S1

We used these formulae to compute the magnetic field $H = B/\mu_0$ within the microchannels between the micromagnets (blue shaded plane in Fig S1A, 0.5 μm above the coverslip surface).

**NdFeB bias magnetic field**: First, we computed the bias magnetic field from NdFeB magnets (*$H_b$* – Eq. S2a). We used 2 NdFeB magnets (1”x1”x1”) placed 1.5–2 cm apart to provide uniform field strong enough to magnetize the array of permalloy micromagents in the MMD. The z-component of the bias magnetic field induction $\mu_0 \vec{H}_{bz}$ near the micromagnets at the midpoint between the two NdFeB magnets is 0.24 T (using the saturation magnetization of NdFeB as $M_s$ ~800,000 A/m). This bias magnetic field can easily magnetize the soft magnetic permalloy micromagnets to saturation (the MMD is saturated at an applied field of ~ 350 A/m i.e 0.44 mT as shown in Fig 2 of main text).

$$\vec{H}_b = \vec{H}_b^{(1)} + \vec{H}_b^{(2)}$$ Eq. S2a

**Permalloy Micromagnetic field:** Based on the analysis above the permalloy micromagnets are driven to saturation by the bias field. Therefore we treated them as independent field sources (permanent bar magnets) with fixed magnetization $M_s$ ~ 860,000. We then computed the field from the arrays of micromagnets ($H_m$ – Eq. S2b) and the total field within the microchannel ($H_T$ - Eq. S2c). The z-component of total magnetic field induction $\mu_0 H_{Tz}$ within the microchannel is > 0.45 T.

$$\vec{H}_m = \sum_i \vec{H}_m^{(i)} \qquad \text{Eq. S2b}$$

$$\vec{H}_T(x,y,z) = \vec{H}_b + \vec{H}_p \qquad \text{Eq. S2c}$$

**Magnetic field gradients within the microchannels:** We derived analytical expressions based on Eq. S1 to compute the magnetic field gradients. The smoothly varying bias magnetic field contributes little to the magnetic field gradients within the microchannels. Magnetic gradients > $10^5$ T/m were obtained about 0.25 µm from the edge of the micromagnets as shown in Fig S1C. The magnetic force structure within the microchannels closely resembles the magnetic gradient profiles (see below).

#### *2.2 Magnetic forces on MNPs - Linear magnetization model:*

We then computed the magnetic forces on MNPs within the microchannels of the MMDs using the linear magnetization model reported earlier[5,7]. Briefly, the magnetization of the MNP (superparamagnetic iron oxide nanoparticle) is assumed to be a linear function of the applied field up to saturation ($M_{ps}$) and a constant thereafter. The equation for the magnetic force on the MNP due to the magnetic field $H_T$ is derived earlier[5] and is given by Eq. S3.

$$\vec{F}_m(x,y,z) = \mu_0 V_p f(H_T)\left(\vec{H}_T \bullet \nabla\right)\vec{H}_T; \qquad H_T = \left|\vec{H}_T\right|$$

$$f(H_T) = \frac{3(\chi_p - \chi_f)}{(\chi_p - \chi_f)+3} \quad for \quad H_T < \frac{(\chi_p - \chi_f)+3}{3\chi_p} M_{ps}$$

$$f(H_T) = \frac{M_{ps}}{H_T} \quad for \quad H_T \geq \frac{(\chi_p - \chi_f)+3}{3\chi_p} M_{ps} \qquad \text{Eq. S3}$$

where $V_p$ is the volume of particle, $M_{ps}$~313,000 A/m is the saturation magnetization of the particle, $\chi_p = (\mu_p/\mu_0 - 1)$ and $\chi_f = (\mu_f/\mu_0 - 1)$ are the magnetic susceptibilities of the particle and fluid respectively; $\mu_p$, $\mu_f$ are the permeabilities of the particle and fluid respectively; $\mu_0$ is the permeability of free space. The formulae for the magnetic field from Eq. S2c and Eq. S1 are used in Eq. S3 to derive the analytical expressions for the 3D magnetic force $F_m(x,y,z)$ analogous to the 2D equations reported earlier[5].

Fig. S2A shows the directionality of the force structure within the microchannel between two micromagnets in the MMD (in the *y-z* plane for a fixed value of $x$ = 0.5µm above the surface of coverslip, since the MNPs within axons are typically confined around this surface). Fig. S2B,C show the dominant force components within the microchannel (along the dotted red line in Fig S2A) and outside the microchannel (along the dotted green, blue lines in Fig S2A). Within the microchannel, the force is predominantly $F_z$ along the z-axis pulling the MNPs to the edge of the micromagnets. While the center of the microchannel is a zero-force zone (zero-gradient), we computed a magnetic force $F_z$ =6.8 pN on a 100 nm MNP 2.5 µm away from the edge of magnet within the microchannel. For MNPs outside the channels, the force is predominantly $F_y$ along the y-axis acting inward towards the channel exits. At a distance of 3 µm from the channel exit, the force acting inward to the channels on a 100nm MNP is $F_y$ =2pN. As the MNPs get closer to the channel exits, a combination of $F_z$, $F_y$ pull the MNPs towards the corners of the magnets. Similar force structure was reported earlier by Gassner *et al.*[8], who used finite element simulations to study the magnetic forces produced by rectangular permanent magnets in a 2D microsystem analogous to our MMDs.

### 3. Supplementary data and processing methods

##### ***3.1 Mapping between MNP Size (SEM diameter) and MNP Fluorescence:***

The MNPs used in this study with streptavidin surface functionalization (binding 7000 pmol biotin/mg) are coated with a high density of dye molecules proportional to the MNP surface area. To the extent that the 561nm laser absorption by iron oxide and self-quenching of Atto-565 are not major factors, we expected a near-linear correlation between the MNP surface area and the MNP fluorescence signal. Fig. S3A shows the square root of fluorescence intensity of MNPs immobilized on coverslips and imaged in our oblique illumination imaging setup. We also overlaid the SEM diameter distribution in Fig. S3A (top axis), which shows a clear mapping between the MNP diameter $D$ and the MNP fluorescence signal $\sqrt{I}$. By using a simple linear mapping ($D=A*\sqrt{I}+B$) we observed good overlap between the cumulative distribution functions of $D$ and $\sqrt{I}$ up to the MNP size of 300 nm as shown in Fig. S3B. This mapping served as a good basis in this work to estimate the MNP diameter given the MNP fluorescence intensity.

##### ***3.2 Magnetic force estimation in glycerol-water mixtures:***

In order to estimate the magnetic forces on MNPs within the microchannels of SMDs we recorded the trajectories of MNPs under force in 10:1 glycerol mixtures as follows. A 10-20μl drop of MNPs in the glycerol:water mixture was placed on top of the micromagnets (i.e. inside and outside the microchannels). After adding the drop, we gently placed a small 4mm x 4mm PDMS slab on top of the drop to minimize any effects of air currents and evaporation. A thin film of MNP solution still exists on top of the micromagnets. Once the particle flow stabilized (monitoring MNP fluorescence in the oblique illumination imaging setup), we suddenly introduced the NdFeB bias magnetic field to magnetize the permalloy micromagnets and thereby turned the magnetic forces on within the microchannels. Simultaneously, we captured the trajectories of MNPs pulled towards the edges of micromagnets by the magnetic forces by imaging the MNPs at 145 fps (Movie S1). When the magnetic force is turned on, we can see the MNPs being pulled into the microchannels in accordance with the direction of the magnetic forces. While the MNPs residing outside the microchannels are pulled inwards parallel to the channels, the MNPs showing up inside the channels are being pulled from the top (Z-axis) towards the middle of the channels.

We used the kymograph processing as described earlier[3] to extract the MNP trajectories from the time lapse movies. Briefly, the time-projection of a movie (Fig. S4A) shows the positional tracks of MNPs that are pulled to the edge of micromagnets from the surroundings. For each track seen in the time-projection, we manually traced the track (Fig. S4A in red) till the edge of the micromagnet to generate a kymograph depicting the intensity variation (due to MNP motion) along the track (Fig. S4B). We then performed 1D-Gaussian fitting of the intensity profile at each time point[3] to extract the particle trajectory *q(t),* where *q* is the distance of the MNP from the edge of magnet (a few sample trajectories shown in Fig. S4C in blue). We then analyzed the MNP trajectories in two different categories, A) within the microchannel and B) outside the microchannels at the channel exits, in order to estimate the forces on MNPs within and outside the microchannels.

Within microchannels: We fit a smooth polynomial to the MNP trajectory *q(t)* as shown in Fig. S4C (in red) and extracted the MNP velocity $v(q=2.5\ \mu m)$ at a distance of 2.5 μm from the micromagnet. We then estimated the diameter of the MNP from the fluorescence intensity as described in the previous section. Using the Stokes relation ($F = 3\pi\eta vD$) we then obtained the force $F(q=2.5\ \mu m)$ exerted on the MNP in the MMD. By processing multiple MNP trajectories we were able to obtain the *F vs D* plot shown in Fig. 3 of main text.

Outside microchannels: In this case (MNPs outside channels approaching the channel exits by moving parallel to the channels before getting trapped by the magnet edge), we computed the $F(q= 3\mu m)$ as a function of MNP diameter, where q is the distance from the channel exit. This data is shown in Fig. 3 of main text.

##### *3.3 Time-lapse movie processing to generate MNP-endosome trajectories:*

In this work we used two different approaches to extract the MNP-endosome trajectories from the time-lapse movies. In the case where the 2D positional track of MNP-endosome *x(t), y(t)* was required, we used the single-particle tracking approach described by us previously[4]. This is the case for the data shown in Figs 5C, 6 and 7. In the case where the 1D motion of MNP-endosome *q(t)* along the microtubule was sought (for velocity estimation), we used the kymograph processing described by us recently elsewhere[3]. This is the case for the data shown in Fig. 4C.


### Supplementary Information: Figures

#### Figure S1:

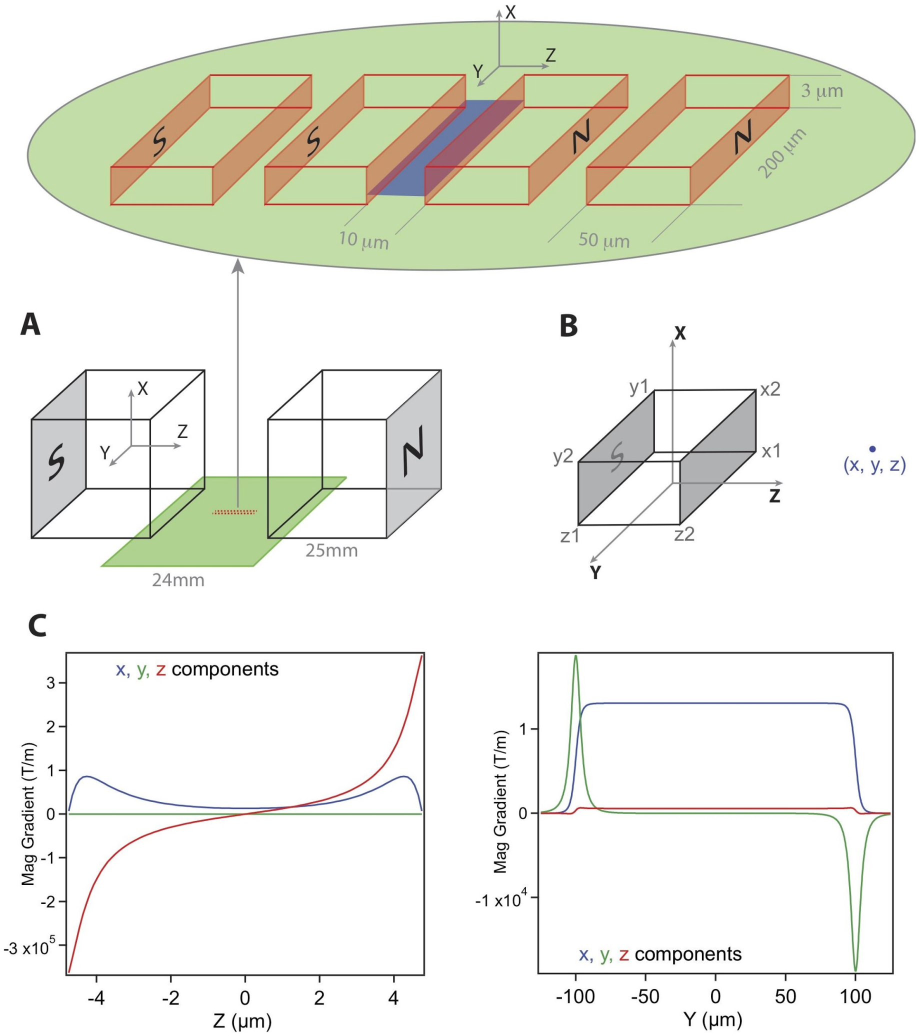


Figure S1: Magnetic manipulation setup and field polarization geometry. A) NdFeB magnets (1" cubes) spaced 1.5-2cm apart provide the bias magnetic field along the Z-axis to magnetize the arrays of permalloy micromagnets (red) on coverslip surface (green). The zoom detail shows a segment of one of the arrays of micromagnets (red) and the surface of interest (blue plane) within the microchannel where the axons carrying the MNPs are confined. B) The edge coordinates of a rectangular bar magnet specified in Eq. S1. C) The *x, y, z* components of the magnetic gradient along the axes bisecting the blue plane (in Fig S1A) with origin at the center of the plane.

### Figure S2:

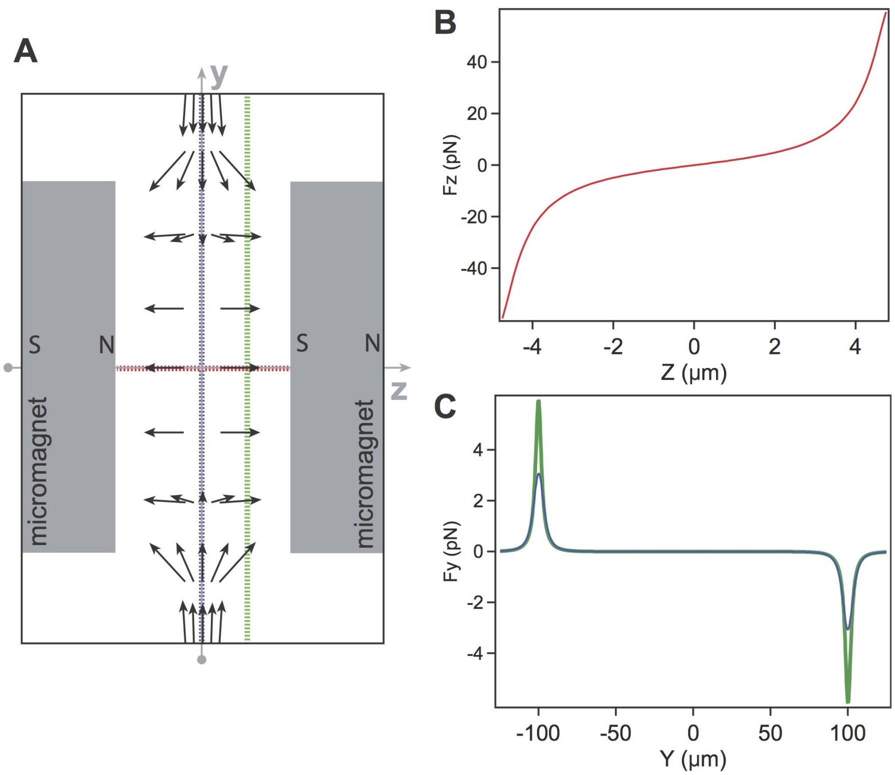


Figure S2: Magnetic force structure between two micromagnets. A) Magnetic force directionality on the blue plane shown in Fig S1A. B) Force profile within the channel along the z-axis (on the red dotted line in A). C) Force profile along the y-axis (on the blue/green dotted lines in A).

### Figure S3:

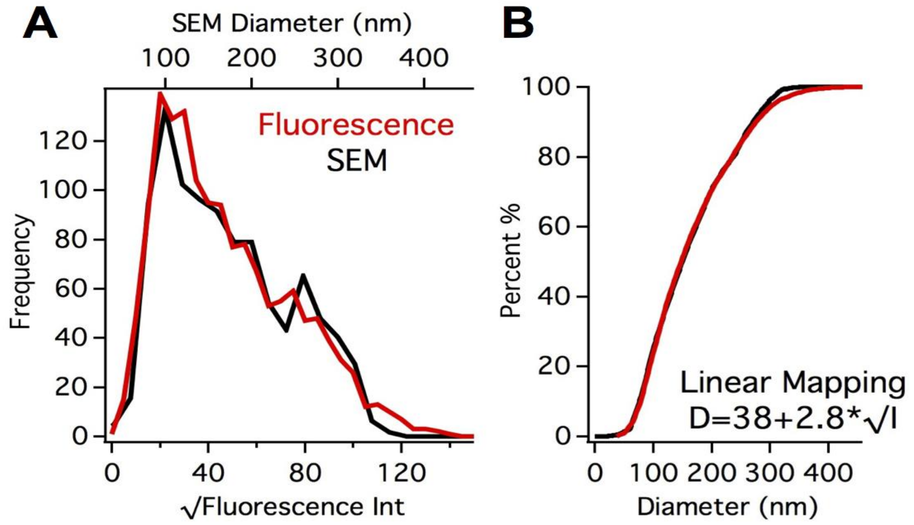


Figure S3: MNP Size and MNP Fluorescence Intensity. A) Distributions of the square root of fluorescence Intensity of MNPs ($\sqrt{I}$, bottom axis, black) and the SEM diameter of MNPs ($D_{sem}$, top axis, red). B) Cumulative distribution functions of $D_{sem}$ (black) and $D_{Fluor}$ (red), where $D_{Fluor}$ is the diameter mapped from the fluorescence intensity of MNP using $D=A+B*\sqrt{I}$.

### Figure S4:

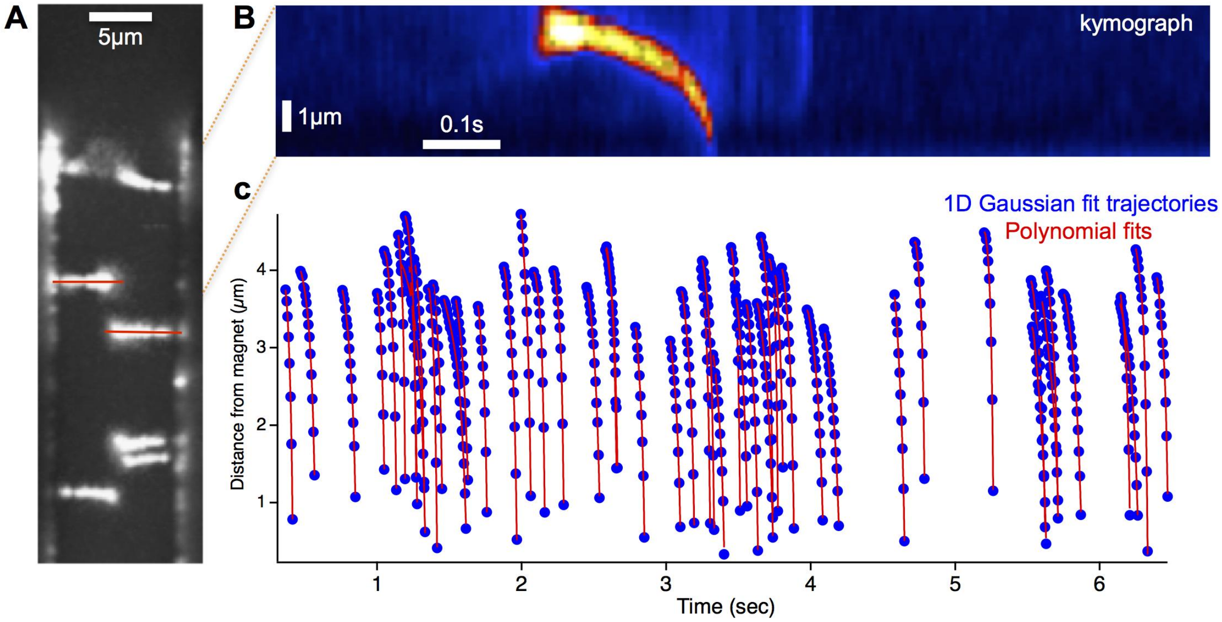


Figure S4: Processing of MNP trajectories in 10:1 glycerol:water mixtures. A) Time projection of a time-lapse movie showing the tracks of MNPs under force being captured on the edges of micromagnets. The red lines are manually traced MNP tracks. B) The kymograph corresponding to a track in (A) depicting the intensity variation along the track due to MNP motion towards the magnet. C) A few sample MNP trajectories (blue markers) obtained by 1D Gaussian fitting of multiple kymographs (from multiple MNP tracks). Also shown in red are the polynomial fits to the trajectories, which are then used to extract the MNP velocities at a given distance from the edge of magnets.

#### Supporting Information: Movie Captions

**Movie S1:** MNP trajectories under magnetic force in glycerol:water mixture in the magnetic-microfluidic device.

**Movie S2:** Retrograde transport of MNP-endosomes in axons in the magnetic-microfluidic device. Imaging at 6.7 fps under 561nm illumination at 0.14 $W/cm^2$. Movie is played at 5x the original speed.

**Movie S3:** Capture of a retrograde MNP-endosome in axon at the edge of the micromagnet. Imaging at 6.7 fps under 561 nm illumination at 0.14 $W/cm^2$. Movie is played at 5x the original speed. Estimated MNP size ~255 nm.

**Movie S4:** Multiple capture and release cycles of a retrograde MNP-endosome in axon at the edge of the micromagnet. Imaging at 6.7 fps under 561 nm illumination at 0.14 $W/cm^2$. Movie is played at 5x the original speed. Estimated MNP size ~259 nm.

**Movie S5:** Capture and release of a retrograde MNP-endosome in axon at the edge of the micromagnet. Imaging at 6.7 fps under 561 nm illumination at 0.14 $W/cm^2$. Movie is played at 5x the original speed. Estimated MNP size ~148 nm.

**Movie S6:** Magnetic force perturbation of retrograde MNP-endosome transport at the channel exit in the magnetic-microfluidic device. Imaging at 6.7 fps under 561 nm illumination at 0.14 $W/cm^2$. Movie is played at 5x the original speed.
